# Depletion and recovery of IgG following treatment with Rozanoliximab and Imlifidase in pigtail macaques

**DOI:** 10.1101/2025.11.11.687913

**Authors:** Kara M. Rzasa, Joanna Zikos, Toni Penney, Faith R. Schiro, Lara A. Doyle-Meyers, Pyone P. Aye, Ronald S. Veazey, James A. Hoxie, Nicholas J. Maness, Diogo M. Magnani, Margaret E. Ackerman

## Abstract

Antibodies are central players in adaptive immunity, providing protection against a wide array of pathogens through mechanisms such as neutralization, opsonization, recruitment of effector immune cells, complement activation and engagement. However, in other contexts, these same effector functions can contribute to immunopathology, particularly when antibodies are developed against self-antigens, resulting in autoimmunity. Understanding the role antibodies play in preventing or causing disease is often supported by studies in model systems wherein manipulation of IgG levels can be used as an experimental tool. Here, we report in simian immunodeficiency virus (SIV) infected pigtail macaques (*Macaca nemestrina*) the capacity of two orthogonal strategies to systemically deplete IgG – treatment with a neonatal Fc receptor blocking antibody (Rozanolixizumab) that restricts IgG rescue and recycling, and administration of the IgG protease Imlifidase (IdeS) that cleaves the Fc domain. Under the conditions evaluated, we observed more rapid and effective, although not necessarily more durable, IgG depletion mediated by IdeS, reducing levels by 74.1-95.1%, compared to a lesser reduction of 31.3-66.9% with anti-FcRn treatment. We observed a similar degree of depletion, comparable kinetics of rebound among SIV antigen-specific fractions as total IgG, but differential balance among IgG subclasses following treatment in some cases. In sum, this study in a nonhuman primate model describes the efficacy and downstream impacts of new tools to modify humoral immune states providing insight into the balance between protective and pathological effects of IgG antibodies.

## Introduction

Antibodies contribute to host protection not only by directly neutralizing pathogens but also by engaging Fc receptors and complement-dependent effector functions to eliminate infected cells^1–3^. Among the antibody isotypes, immunoglobulin G (IgG) represents the most abundant class in circulation, accounting for approximately 70–85% of total serum immunoglobulins^1–3^. IgG is further divided into four subclasses IgG1, IgG2, IgG3, and IgG4 named in decreasing order of relative abundance^1–3^. In humans, these subclasses are both structurally and functionally distinct, based on differences in their hinge regions, affinity for Fcy receptors (FcyR), and capacity to activate complement^2^. IgG1 and IgG3 are generally the most potent in activating the classical complement pathway, mediating antibody-dependent cellular cytotoxicity (ADCC) and antibody-dependent cellular phagocytosis (ADCP) through Fcγ receptor engagement^2–5^. IgG2 is less efficient at these functions and considered to have a lower inflammatory profile but is particularly important for responses against carbohydrate antigens like bacterial polysaccharides^2,6,7^. IgG4 also exhibits minimal ability to activate complement, is functionally monovalent due to Fab-arm exchange, and generally demonstrates anti-inflammatory properties, although it can still contribute to pathology in certain IgG4-related diseases^1–3,8,9^. Functional differences among IgG subclasses are less pronounced in non-human primates^10–13^, but nonetheless offer a means to diversify the activity and function of IgG in circulation and transport to protect mucosal sites.

The dominant IgG Fc mechanisms include complement activation and deposition, ADCC, and ADCP^1–3,5^. These same effector pathways, however, are co-opted by pathogenic autoantibodies that cause autoimmune disease, in which they drive various inflammatory pathways leading to tissue injury and organ dysfunction^14–17^. Many pathogenic autoreactive IgG responses are mediated through excessive complement activation, leading to target cell lysis, and recruitment and activation of innate effector cells driving local inflammation^15,16^.

One approach to interrogate the role of IgG antibodies *in vivo* is by determining if selective IgG depletion modulates disease activity in the host^18–22^. In individuals with autoimmune disease driven by pathogenic IgG, depletion often correlates with clinical improvement or disease reversal^19–21,23^. Conversely, in settings in which IgG antibodies are required for protective immunity, such as containment of latent viral infections or control of extracellular bacteria, antibody depletion can impair immune surveillance and control, resulting in pathogen reactivation and accelerated disease progression^24–27^. Thus, given that IgG effector functions range from contributing to protective immunity to pathogenic autoimmunity, there is great interest in approaches to modulate IgG levels *in vivo* to understand their physiologic functions and pathophysiologic mechanisms and as a therapeutic intervention.

In particular, depletion of IgG antibodies can be useful in the treatment of IgG-mediated autoimmune diseases, IgG-mediated graft rejection or fetal toxicity (alloimmune disease), as well as in studies of IgG-mediated pathogen control^21,26,27^. IgG undergoes recycling and rescue from degradation in the lysosome via the neonatal Fc receptor (FcRn) in a pH-dependent manner. IgG binds to two FcRn molecules at low pH within endosomes, allowing recycling back to the cell surface, where at neutral pH it is released back into circulation^18–20^. FcRn antagonists such as Efgartigimod^28^, Nipocalimab^29^, Batoclimab^23^, Orilanolimab^18,20^, and Rozanolixizumab^30–33^ are antibody-based therapies used clinically to treat various autoimmune conditions. With the exception of Efgartigimod, these treatments work by targeting FcRn through the Fab region of the antibody, which binds the same IgG Fc binding site but across varying pH conditions, thereby blocking IgG recycling^19,20,34^. Efgartigimod is an IgG1 Fc portion that instead binds FcRn in the same pH dependent manner of native IgG, but with higher affinity afforded by incorporation of a set of five amino acid substitutions^28,35^. Thus, these antagonists bind and block FcRn, reducing binding to endogenous IgG, preventing recycling, and resulting in IgG degradation in lysosomes and reducing circulating IgG levels^19,20^. FcRn blockade is temporary as these antibodies/fragments are eventually degraded by the host, allowing FcRn recycling pathways to recover, with IgG levels returning to baseline^18,20,23,28–30^.

In humans, FcRn blocking therapies are under active investigation or are currently in use for treatment of a wide range of antibody-association pathologies. Myasthenia gravis, an autoimmune disease mediated by autoantibodies against the nicotinic acetylcholine receptor, is a disease in which IgG depletion has shown efficacy^36,37^. Other conditions such as thyroid autoimmunity^38^, autoimmune encephalitis, chronic inflammatory demyelinating polyneuropathy, immune thrombocytopenia, bullous pemphigoid, pemphigus, warm autoimmune hemolytic anemia, rheumatoid arthritis, systemic lupus erythematosus, severe fibromyalgia, graves’ ophthalmopathy, and hemolytic disease of the fetus/newborn are currently under clinical evaluation for IgG depletion efficacy by FcRn blockade^39,40^. These promising clinical advances underscore the transformative potential of FcRn blockade as a targeted strategy to rapidly and safely reduce pathogenic IgG autoantibodies across a diverse spectrum of autoimmune and antibody-mediated diseases.

The clinical utility of targeted approaches for IgG depletion has also motivated development of alternative strategies that modify IgG function. Enzymes from various bacteria that specifically cleave the human IgG Fc domain (IdeS^21,22^, SpeB^41^, Gingipain K^42^ and IceMG^43^) or the conserved Fc glycan (EndoS^41,44^) that are required for interaction with Fc receptors and elicit antibody effector functions have been increasingly used in biotechnology applications. This approach recently gained a clinical foothold with the approval of IdeS for treatment of highly sensitized patients awaiting kidney transplantation by rapidly and effectively cleaving circulating IgG antibodies, including pathogenic donor-specific antibodies, thereby preventing antibody-mediated rejection and enabling allo-transplantation in to be performed^21,45^.

Imlifidase (IdeS), an enzyme from *Streptococcus pyogenes* that cleaves the Fc from the Fab domains, separates FcyR-binding from antigen-binding antibody domains^21,22,44,46,47^. IdeS is the only IgG-degrading enzyme approved for clinical use in humans; alternatives used in preclinical models and *in vitro* include SpeB^41^, Gingipain K^42^, and IceMG^43^. IdeS cleaves IgG just below the hinge in two steps: the first cleaves one Fc, leaving single-cleaved IgG (scIgG), and the second cleaves the other Fc, producing Fab and Fc/2 fragments^21,22^. In humans and animal models, IdeS acts quickly, causing a drastic drop in IgG levels within two hours of administration and with recovery occurring within one to two weeks^21,47^.

While many of these approaches to deplete IgG have been investigated in nonhuman primate models^21,41,43^, none have been evaluated in pigtail macaques, which are important models for infectious diseases including influenza, tuberculosis^48^, COVID-19^49^, and non-human primate models of AIDS^50^, as well as in studies of fertility, fetal development^51^, and organ transplantation^52^. Here we evaluate the efficacy of the anti-FcRn antibody Rozanolixizumab and the IgG cleaving enzyme IdeS in depleting IgG in pigtail macaques, and report how levels of major immunoglobulin classes, antigen-specific IgG subclasses, and antibody functions are impacted by these interventions.

## Materials and methods

### Research Animals

Male pigtail macaques (*Macaca nemestrina*) were inoculated intravenously with SIVmac239ΔGY (300 TCID_50_), a variant of simian immunodeficiency virus^53^, as part of an NIH pathogenesis and vaccine study (Simian Vaccine Evaluation Unit P191; Contract number: HHSN2722013000041; Task Order Number: HHSN27200006) followed by intravenous challenges with SIVsmE660 (100 TCID_50_). Blood was collected throughout the study, along with physical examinations and blood collection for monitoring serum chemistry and complete blood counts for clinical evaluation. To assess the role of IgG in viral control, Rozanoliximab was administered under anesthesia to three animals via intravenous infusion at 30 mg/kg over four total doses, each separated by one week. IdeS was administered under anesthesia to these same animals via intravenous administration at 1.5 mg/kg on five consecutive days. The IdeS treatment was repeated in two of these animals.

### Ethics Statement

Pigtail macaques used in this study were purpose bred at the Washington National Primate Research Center and moved to the Tulane National Biomedical Research Center (TNBRC, formerly the Tulane National Primate Research Center) for these experiments. Macaques were housed in compliance with the NRC Guide for the Care and Use of Laboratory Animals and the Animal Welfare Act. Animal experiments were approved by the Institutional Animal Care and Use Committee of Tulane University. The TNBRC is fully accredited by AAALAC International (Association for the Assessment and Accreditation of Laboratory Animal Care), Animal Welfare Assurance No. A3180-01.

### Sequence and structure visualization

Alignments of human, rhesus, cynomolgus, and pigtail macaque FcRn and IgG (**Supplemental Table 1**) were generated using Geneious. The structures of human FcRn bound to Rozanolixuzumab (6fgB) and IdeS bound to human IgG1 Fc (8A47) were retrieved from the Protein Data Bank. The pigtail FcRn sequence was retrieved from Uniprot (A0A2K6CTM0). Structural analyses were performed with UCSF ChimeraX (Resource for Biocomputing, Visualization, and Informatics at the University of California, San Francisco)^54^.

### Rhesus Rozanolixuzumab (Roz)

Primate chimeric anti-FcRn monoclonal antibodies were designed based on the clinically developed anti-FcRn antibody Rozanolixizumab, which binds a conserved FcRn epitope critical for IgG interaction. The antibodies were specifically engineered to inhibit FcRn-mediated recycling of IgG in nonhuman primates. Initially, rozanolixizumab was re-engineered as a ‘primatized’ rhesus-chimeric IgG4 to match the human therapeutic IgG subclass. However, rhesus IgG4 exhibited weaker heavy–light chain pairing due to the atypical absence of cysteine (C131) in the rhesus IgG4 heavy chain, critical for disulfide linkage^55^. To address potentially confounding issues in developability and potency arising from this variant phenotype, we re-engineered the antibody as a rhesus IgG1 isotype incorporating ‘Fc silencing’ LALA mutations to mitigate FcyR activation. This rhesus chimeric antibody, termed anti-FcRn [RozR1LALA], was produced by the Nonhuman Primate Reagent Resource (NHPRR; nhpreagents.org). An independent anti-FcRn clone, M281, expressed as an aglycosylated chimeric rhesus IgG1 with an N297G mutation, was also developed to confirm binding specificity.

The ‘rhesusized’ anti-FcRn monoclonal antibody, Anti-FcRn [RozR1LALA] (NHPRR Catalog, CAT-00095, RRID: AB_2888630, lot PPLOT00189), hereafter referred to as Roz, was produced from a Chinese Hamster Ovary (CHO) stable cell line cultured in a controlled bioreactor system under quality unit oversight to generate safe, effective, and reproducible preclinical-grade material. Post-production, Roz was purified and formulated in a buffer comprising 20 mM sodium citrate, 150 mM sodium chloride (NaCl), and 0.02% polysorbate 80 (PS80), adjusted to pH 6.0. Analytical testing confirmed suitability for infusion, demonstrating binding activity by ELISA, endotoxin levels below 0.1 EU/mg, and a purity exceeding 99% monomer content by size-exclusion chromatograph. Fluorescently-labeled Roz (Anti-FcRn [RozR1LALA]-FITC, NHPRR Cat# CAT-00256, RRID: AB_2941340) and Anti-FcRn [M281]-FITC (NHPRR, CAT-00035, RRID: AB_2941348) and Anti-FcRn [RozR1LALA]-FITC were also generated.

### Roz binding to FcRn

The transiently expressed CHO-derived antibodies were evaluated for binding to soluble FcRn. Binding of FcRn-specific antibodies to Cynomolgus/Rhesus macaque FcRn heterodimer protein (FCGRT&B2M, His Tag & Strep II Tag, SPR verified, ACRO Biosystems Cat# FCM-C5284-50ug) and recombinant soluble human FcRn (PDB:6FGB_A) was defined. Human FcRn was expressed as the extracellular domain co-transfected with human β2-microglobulin (B2M) using VRC 2092 (CMV8xR mIL2L-hFcRn_dTMdL) and VRC 2093 (CMV8xR human B2M-His) using plasmids obtained from VRC/NIAID. Briefly, enzyme-linked immunosorbent assays (ELISA) were conducted using 96-well high-binding polystyrene microplates coated with the purified soluble FcRn. The primary antibodies, transiently produced in CHO cells, were incubated to allow binding to the immobilized FcRn. Specific antibody binding was subsequently detected and quantified using standard ELISA techniques, employing appropriate secondary reagents and colorimetric readouts. Controls included independent anti-FcRn clones to ensure assay specificity and reproducibility.

### Generation and validation of IdeS

A DNA sequence encoding the Immunoglobulin G-degrading enzyme of *Streptococcus pyogenes* (IdeS), Uniprot accession F8V4V0^56^, was expressed using the pET-30a (+) vector. The sequence was codon-optimized for *E. coli* expression, and the construct was fused to a TEV cleavage site followed by an 8X His tag and a Twin-StrepII purification tag at the C-terminus^57^. The enzyme was formulated in PBS (pH 6.6) containing 10% glycerol. The preparation is supplied by NHPRR under catalog number CAT-00341 (RRID:SCR_012986). The enzymatic activity of CAT-00341 has been validated both *in vitro* and *in vivo* as previously described^21^.

For this study, lot PPLOT00167 was provided at a concentration of 0.56 mg/mL. Purity was confirmed by SDS-PAGE (≥90% purity), and molecular mass was verified by LC-MS. Endotoxin levels were <0.9 EU/mg. Samples were stored at −80 °C until use.

### Digestion of IgG with IdeS

IdeS (NHPRR: CAT-00341) was incubated with rhesus recombinant monoclonals IgG1 (RRID:AB_2819330), IgG2 (RRID:AB_2819331), IgG3 (RRID:AB_2819332) and IgG4 (RRID:AB_2819333) or diluted pigtail sera for an hour in a 37°C water bath, as previously described^21^. Digested antibodies were separated under non-reducing conditions by NuPage 4-12% Bis-Tris and stained with Coomassie Blue. PageRuler Unstained Protein Ladder was used to determine band molecular weights. Samples were visualized using a ChemiDoc Imaging system and analyzed using the Image Lab Software (Bio-rad).

### Measurements of total and antigen specific-immunoglobulins

Plasma antibody profiles were characterized by the Fc array assay, as described previously^58,59^. Minor modifications for the measurement of total immunoglobulins were used in that mouse anti-monkey IgG, mouse anti-human IgM, and goat anti-human IgA (**Supplemental Table 2**), each of which recognizes or cross-reacts with their respective pigtail macaque Ig isotypes, were covalently coupled to magnetic microspheres using carbodiimide chemistry (Luminex Corporation)^58,59^. The anti-antibody beads were incubated with diluted pigtail sera then detected with mouse anti-monkey IgG, mouse anti-human IgM, or goat anti-human IgA conjugated with R-phycoerythrin (**Supplemental Table 2**). Antibodies specific for a panel of SIV antigens (**Supplemental Table 3**) similarly coupled to multiplex assay beads were quantified by detection of total immunoglobulins or FcyR binding (**Supplemental Table 3**). Initial pilot experiments were used to determine appropriate dilutions used. The median fluorescent intensity (MFI) of detected antibody was acquired on a FlexMap 3D array reader (Luminex Corporation). Fold-change in IgG levels over time relative to a pre-Roz or pre-IdeS timepoint was calculated for total IgG by averaging the fold-change of the raw MFI for dilutions in the linear range of the standard curve (1:40,000, 1:80,000 and 1:160,000). Data was graphed in GraphPad Prism (Version 10.5.0).

### Antibody dependent complement deposition (ADCD)

Antibody-dependent complement deposition (ADCD) experiments were performed as previously described^60^ with minor modifications. Plasma samples were diluted to 1:100 and 1:5000 and incubated for two hours with the antigen-coupled beads (**Supplemental Table 3**). Subsequently, samples were incubated with 1:100 dilution of low tox guinea pig complement (Cedarlane, CL4051) for one hour at 37°C and then another hour with PE-labeled anti-complement C3b antibody (BioLegend, 846103) to measure complement deposition. The median fluorescent intensity (MFI) of detected antibody was acquired on a FlexMap 3D array reader (Luminex Corporation). Data was graphed in GraphPad Prism (Version 10.5.0).

### Statistical Analysis

Data analysis was performed in GraphPad Prism (Version 10.5.0). A two-tailed exact Mann–Whitney U test was used to compare depletion levels between IdeS treatment (n = 7) and Roz treatment (n = 3) due to the small sample sizes and non-normal data distributions; median percentage depletion was determined for each group, and the resulting U statistic (U = 0) and exact p-value (p = 0.0167) indicated a statistically significant difference at p < 0.05, while the Hodges–Lehmann estimator was calculated to provide the median difference between groups (32.6 percentage points), with all analyses conducted according to standard non-parametric statistical guidelines.

## Results

### Structural and sequence homology of human and NHP FcRn

Rozanolixizumab binds to human FcRn at an epitope critical for the interaction of this receptor with endogenous IgG. Alignments of human, rhesus, cynomolgus and pigtail FcRn show a high degree of amino acid sequence conservation between species (**Figure 1A**). Pairwise identity between human and pigtail FcRn is 96.8%, and contact residues identified from the co-crystal structure of Rozanolixizumab with human FcRn^32,33^ are perfectly conserved. While there are several sequence differences between pigtail macaque and humans, apart from one substitution within the FcRn intracellular domain, these changes were conserved with other NHP species in which Rozanolixizumab has shown effective antibody depletion^61^.

**Figure 1:**
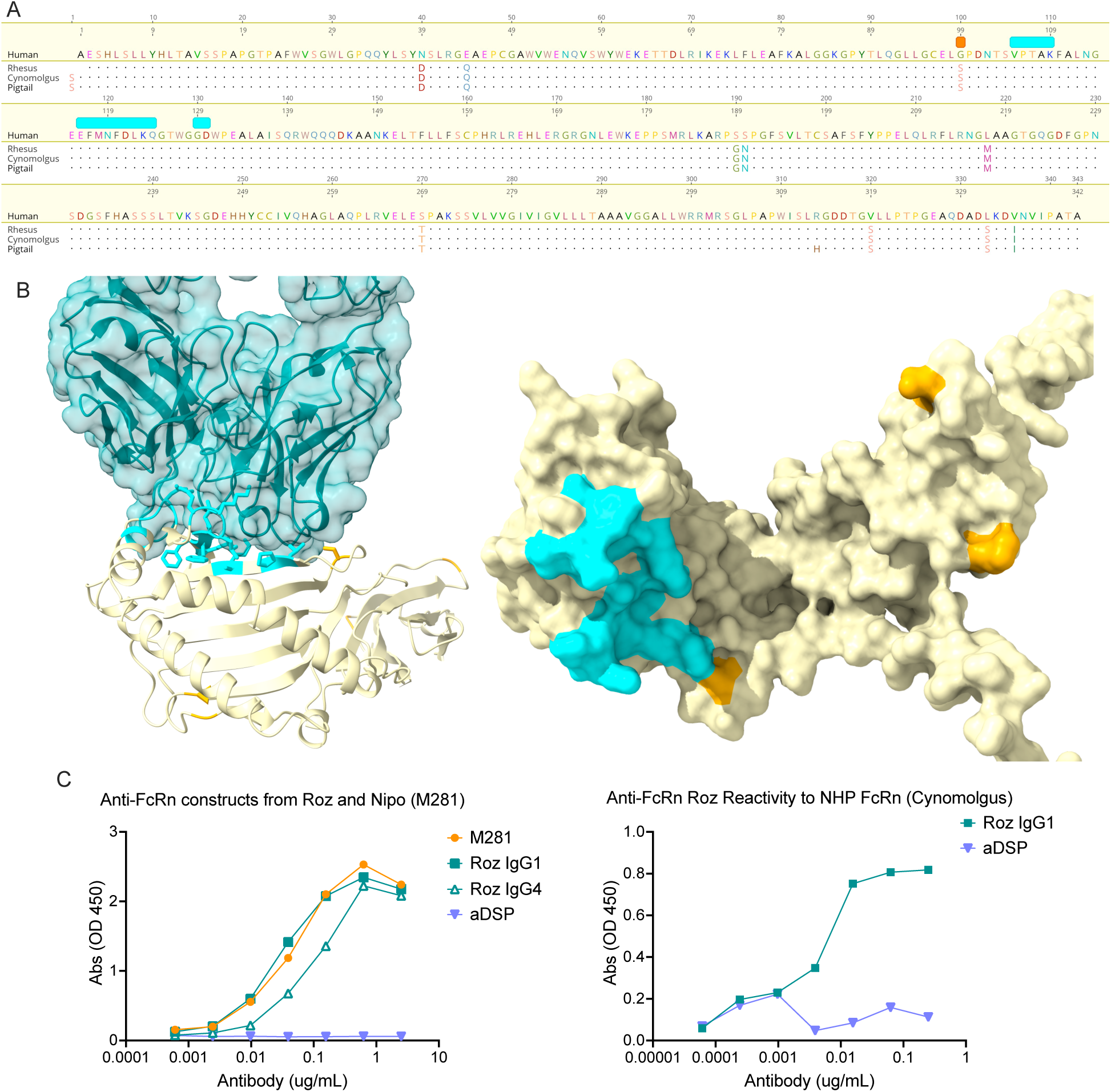
Conserved FcRn sequence, structure and binding of Roz *in vitro*. **A.** Amino acid sequence alignment of human, rhesus, cynomolgus), and pigtail macaque FcRn with reported Roz contact residues in cyan. **B.** (Left) Model of pigtail FcRn (khaki, Uniprot: A0A2K6CTM0) bound to Roz (teal, PDB:6fgB) based on its co-crystal structure complex with chain A of human FcRn (hidden). (Right) surface model of the Roz binding site on FcRn. Amino acids that vary between human and pigtail macaque FcRn are illustrated in orange and contact interface residues (≤3 A) highlighted in cyan. **C.** Binding of Roz to human (left) and rhesus macaque (right) FcRn.

Structural visualization of the sites of sequence variation demonstrates that, except for G99S, these substitutions are distal to the Rozanolixizumab contact region (**Figure 1B**). Collectively, the high degree of sequence and putative structural homology motivated development of a primatized form of Rozanolixizumab for use in these valuable model systems with reduced likelihood of induction of anti-human IgG antibody responses.

### Primatization of Rozanoliximab

‘Primatized’ versions of Rozanoliximab–i.e., Roz antibodies with higher sequence identity to NHP molecules– were specifically engineered to inhibit FcRn-mediated recycling of IgG in NHP. The parental Roz antibody was first expressed as a rhesus-chimeric IgG4 via a ‘constant-region replacement’, in which the human constant domains were swapped for the corresponding rhesus sequences. Although IgG subclass functions are imperfectly conserved between species^62^, the rhesus-chimeric IgG4 nominally matched the human therapeutic IgG subclass, selected for its reduced effector function. However, rhesus IgG4 lacks the heavy chain cysteine (C131) that allows for the covalent interchain disulfide bond linking the heavy and light chains^55^. Consequently, in intact rhesus IgG4, the heavy and light chains rely on noncovalent interactions to maintain heavy and light chain assembly^63^. To mitigate potential developability and potency issues arising from this variant phenotype, we re-engineered the antibody as a rhesus IgG1 isotype incorporating ‘Fc silencing’ LALA mutation^64^ to reduce FcγR activation. This engineered design preserves the reduced effector function potential while improving molecular stability. The resulting rhesus anti-FcRn chimeric antibody [RozR1LALA], herein referred to as Roz, and an independent anti-FcRn clone, M281^65^, expressed as an aglycosylated (N297G) chimeric rhesus IgG1, were developed and used to confirm binding specificity. Binding of both Roz, the rhesus IgG4 variant Roz IgG4, and M281 antibodies to rhesus and human FcRn was confirmed by ELISA (**Figure 1C**).

### Roz function is conserved in pigtail macaques

The ability of Roz to deplete systemic IgG was evaluated in three pigtail macaques (RI-01, RI-02, and RI-03) administered a 30 mg/kg dose by intravenous (IV) infusion, a dose and route that has effectively reduced plasma IgG in prior studies in rhesus and cynomolgus macaques^33,61,66^. To prolong the period of depletion, Roz was administered on days 0, 7, 14 and 21. Plasma samples were collected over time to measure the effect of this treatment on antibody levels. Limited reduction in total plasma IgG was apparent from SDS-PAGE analysis (**Figure 2A**), though some differences were observed among animals (**Supplemental Figure 1**). To quantify the extent and isotype specificity of systemic depletion, total plasma immunoglobulins were measured by multiplex assay. Whereas decreased levels of IgG were apparent at timepoints proximal to Roz administration, no such effect was observed for IgA or IgM (**Figure 2B, Supplemental Figure 2**). Levels of total IgG were quantified as fold-change relative to a pre-treatment timepoint (**Figure 2C**). Depletion varied in extent and kinetic profile, with animals presenting maximal depletion at 7-14 days and plasma IgG levels reduced to between 31.3 – 66.9% [Mean 48.4%, SD± 17.8%] of the level observed pre-treatment. Durability also varied, with one animal returning to baseline levels before the final Roz dose, and another showing near maximal depletion out to one month post the first Roz dose (one week post the last dose). Overall, the somewhat limited degree and durability of IgG depletion that resulted from this FcRn-blocking regimen motivated exploration of alternative strategies.

**Figure 2.**
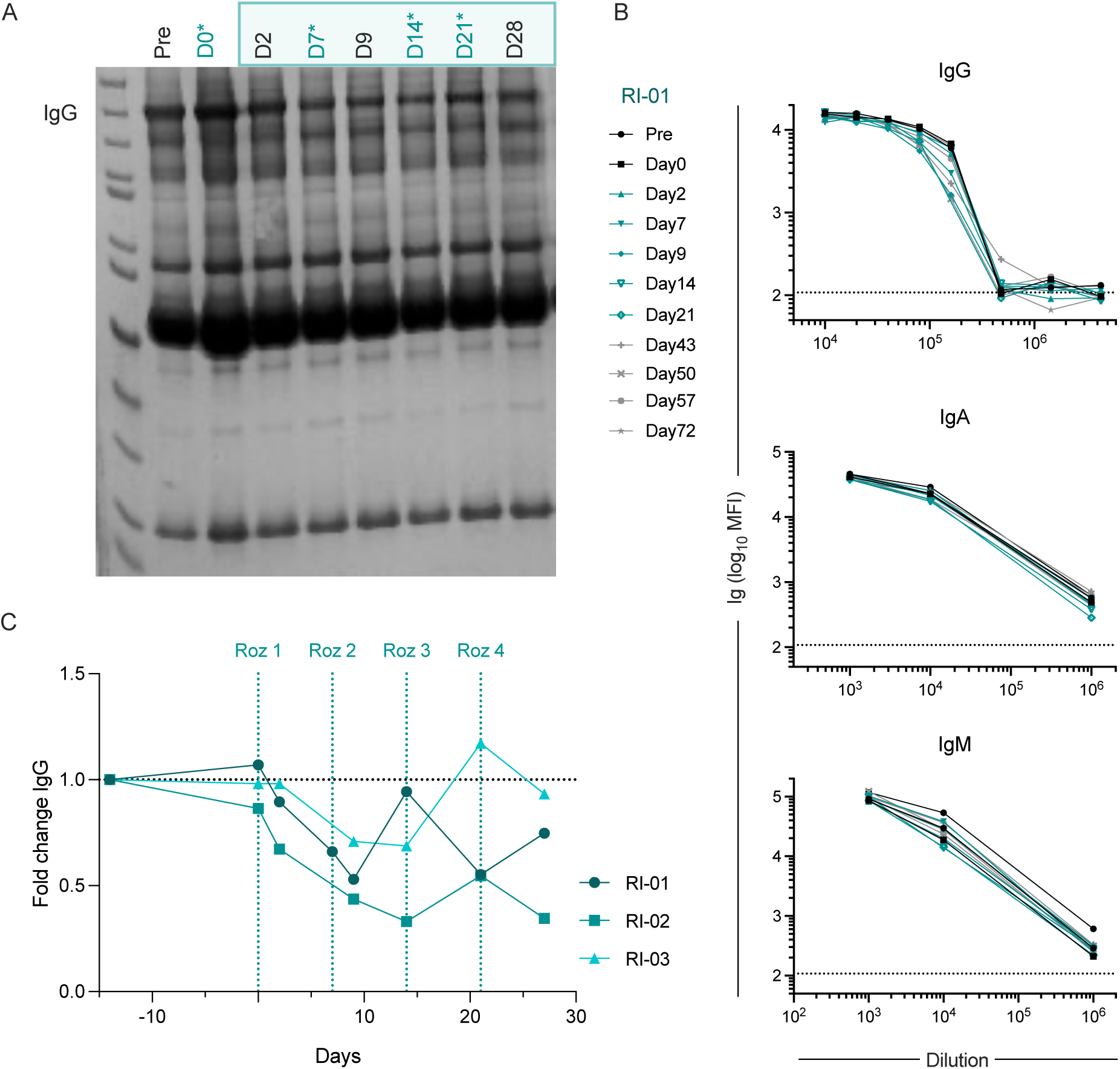
*In vivo* efficacy and specificity of Roz treatment in pigtail macaques. **A.** Representative SDS-PAGE of total serum protein in animal RI-01 over time before, during, and after treatment with Roz. Days on which Roz was administered are indicated with an asterisk and those anticipated to show IgG depletion in the teal box. **B.** Median fluorescent intensity (MFI) of total IgG (top), IgA (center), and IgM (bottom) measured in diluted serum over time in animal RI-01. Color and symbol indicate sample timepoint. Dotted line indicates lower limit of detection. **C.** Fold-change in total serum IgG over time in three animals treated with Roz at indicated timepoints.

### Structural and sequence homology of human and NHP IgG

Restriction of the IgG Fc domain with IdeS represents an alternative strategy to eliminate the FcRn-mediated recycling of IgG that drives the high systemic half-life of IgG molecules. The four IgG subclasses present in humans, rhesus, cynomolgus and pigtail macaques were aligned from the lower hinge through CH2 and CH3 domains (**Figure 3A**). Despite sequence diversity in the cleavage site between G118 and G119, and among contact residues defined from the co-crystal structure of IdeS with human IgG1^67^, IdeS cleaves all human IgG subclasses. Digestion of human IgG2, however, which, among other sequence differences in the lower hinge, bears an AG rather than GG motif at the site of cleavage, is less efficient^22^. In general, the subclasses of all three NHP species show more sequence and functional similarity to each other and to human IgG1 than to other human subclasses^62^. The IdeS binding and cleavage site residues are generally well-conserved, with most of the substitutions between human and pigtail IgG1 located distal to the IdeS binding locations (**Figure 3B**); however, some substitutions, most of which are conserved among NHP and subclasses, occur in the binding site.

**Figure 3.**
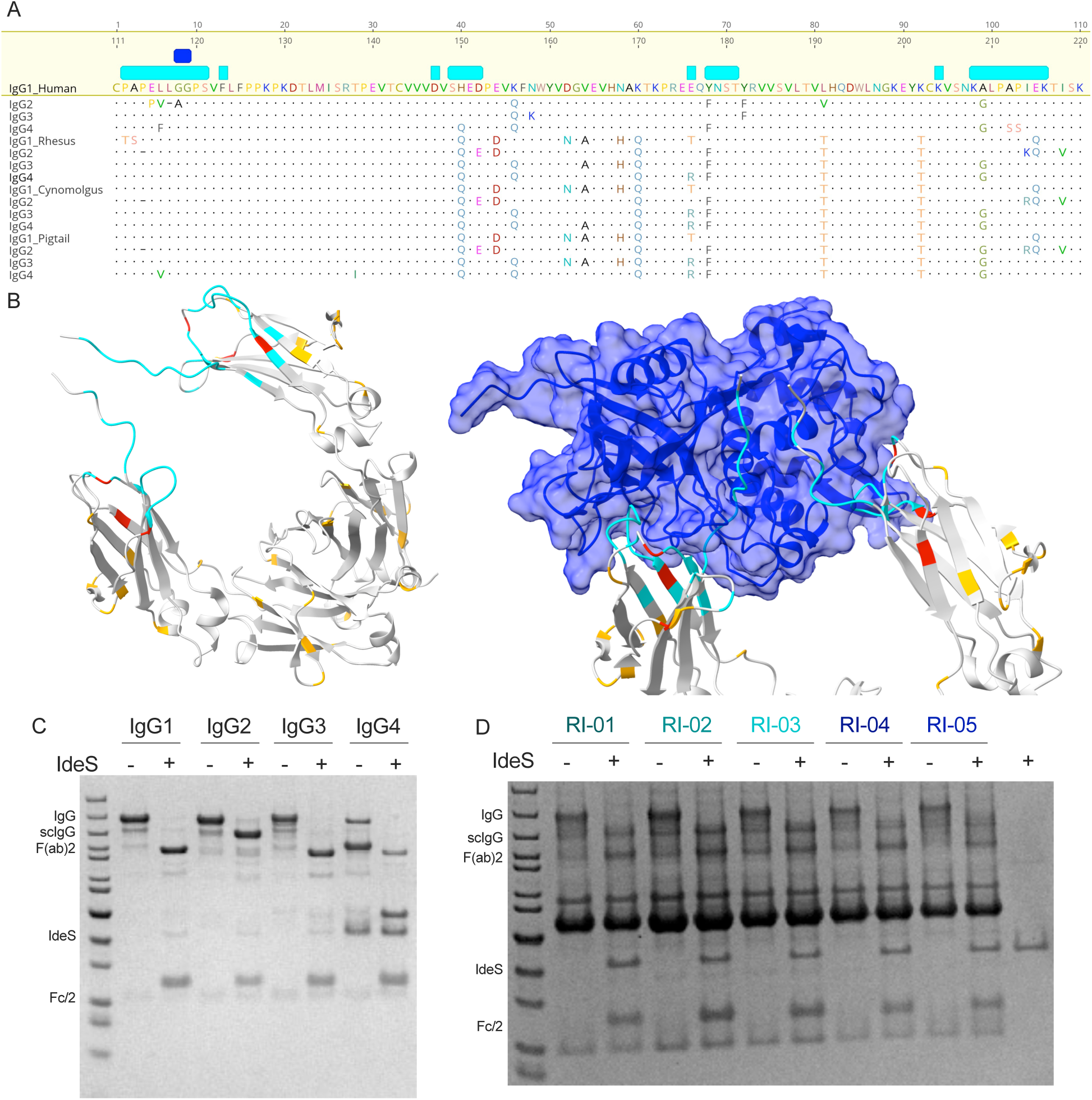
Conserved IgG sequence, structure, and cleavage by IdeS *in vitro*. **A.** Amino acid sequence alignment of human, rhesus, cynomolgus, and pigtail macaque IgG subclasses. IdeS contact residues are indicated in cyan and the IdeS cleavage site in dark blue. **B.** (Left) IgG1 Fc structure. (Right) Co-crystal structure of IdeS (blue) and human IgG1 Fc (silver) (PDB: 8A47). The IdeS binding site (≤3 A) is denoted in cyan. Amino acids that vary between human IgG1 and pigtail macaque IgG1 within (red) and outside (orange) the binding site are indicated. **C-D.** SDS-PAGE of recombinant rhesus IgG subclasses (**C**) and pigtail macaque serum from individual animals (**D**) with (+) or without (-) incubation *in vitro* with IdeS.

### Cleavage of recombinant and plasma-derived NHP IgG in vitro

Though not identical to the sequences in pigtail macaque IgG, recombinant rhesus IgG subclasses were incubated with IdeS and analyzed by SDS-PAGE to evaluate susceptibility to cleavage (**Figure 3C**). All four subclasses showed evidence of cleavage, though rhesus IgG2 was recalcitrant to full digestion, exhibiting a predominant digestion product that lacked one heavy chain CH2 and CH3 domain, representing singly cleaved scIgG. Unlike human IgG2, rhesus IgG2 does not have altered cleavage site amino acids but does bear a deletion of P114, three residues upstream. The conservation of both this deletion as well as D152E and I214K(rhesus)/R(cynomolgus and pigtail) substitutions in putative contact residues in cynomolgus and pigtail macaques suggests the possibility that IdeS also exhibits reduced activity against IgG2 in these species. While other differences also exist between well-cleaved human IgG1 and rhesus IgG2 (e.g. H150Q, E215Q), these substitutions are conserved with rhesus IgG1, which this and prior testing shows is also well-cleaved^21^. Consistent with expectations based on these results and sequence homology, when plasma from each of five pigtail macaques was incubated with IdeS *in vitro*, cleavage was observed (**Figure 3D**). While there was limited evidence of intact IgG, most plasma samples presented apparently similar levels of doubly cleaved and scIgG, consistent with incomplete digestion under the conditions tested, and the reduced efficiency of IdeS in restricting sc as opposed to intact IgG.

### Cleavage of pigtail macaque IgG in vivo

To test the ability of IdeS treatment to deplete plasma IgG in pigtail macaques, enzyme was administered to five animals at a dose of 1.5 mg/kg by IV on each of five consecutive days. Two animals (RI-04 and RI-05) underwent two such rounds of IdeS treatment 101 days apart, and three animals (RI-01, RI-02, and RI-03) were treated with IdeS six days after the last dose of Roz. Plasma samples were collected longitudinally to investigate the effect of IdeS on circulating immunoglobulin levels. Analysis of plasma proteins by SDS-PAGE demonstrated reduction in intact IgG and accumulation of cleavage fragments, including free CH2-CH3 domains (Fc/2), during IdeS treatment (**Figure 4A**, **Supplemental Figure 1**). Whereas neither IgM nor IgA levels changed over time, IgG levels showed considerable depletion when quantified by multiplex assay (**Figure 4B**, **Supplemental Figure 3**). Plasma IgG levels were reduced by 74.1-95.1%, and efficiency was similar for the first regimen as for the second or when treatment followed Roz (**Figure 4C**). Depletion was both effective (Mean 81.9%, SD± 7.41%) and rapid, with reduction in intact IgG observed as early as one day after the first dose and maximal depletion observed within one week. Durability varied, with intact plasma IgG levels returning to pre-treatment levels by eight days post first dose for the animal exhibiting the most rapid recovery, as compared to over one month for the slowest.

**Figure 4.**
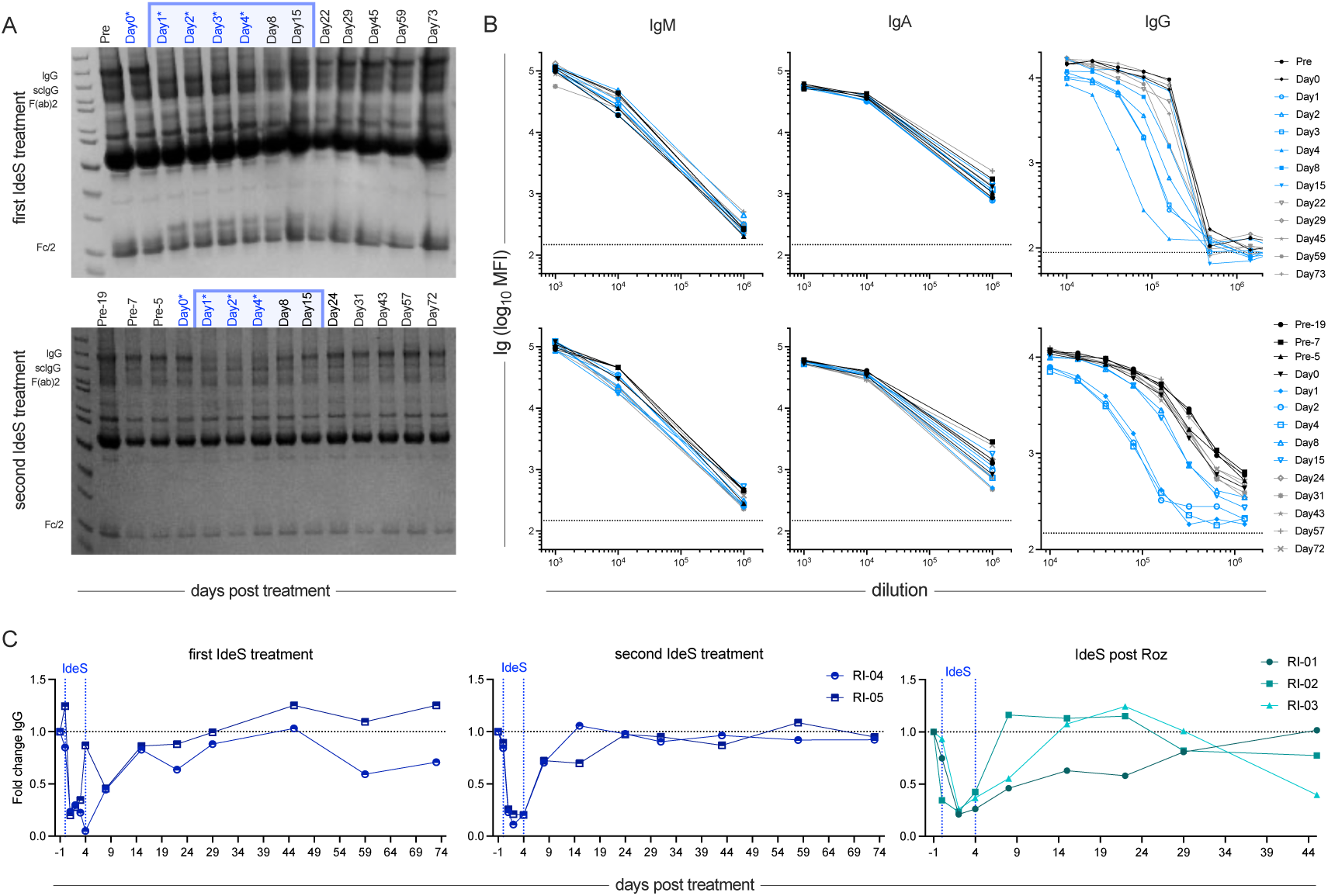
*In vivo* efficacy and specificity of IdeS treatment in pigtail macaques. **A.** Representative SDS-PAGE of total serum protein in animal RI-05 over time during two sequential IdeS treatment periods. Days of IdeS treatment are indicated with an asterisk and those anticipated to show IgG depletion in the blue box. **B.** Median fluorescent intensity (MFI) of total IgM (left), IgA (center), and IgG (right) measured in diluted serum over time. Color and symbol indicate sample timepoint. Dotted line indicates lower limit of detection. **C.** Fold-change in total serum IgG over time in two animals treated with an initial (left) and second (center) IdeS regimen, and in three animals treated following Roz (right). IdeS treatment window (days 0-4) is marked with blue dotted lines.

### Depletion of antigen-specific IgG

Because these experiments were performed in animals infected with SIV, we had the opportunity to evaluate the ability of each intervention to reduce antigen-specific antibodies systemically. The level of SIV-specific IgG over time in plasma were also measured at baseline (sample 1) and following infection (samples 2 and following). Testing across a panel of antigens including envelope (gp120, gp130, gp140), capsid (p17, p27, gag, pr55), and other proteins (rev, rt, tat, nef) originating from SIV239, SIVmac251, or SIVsmE660 provided the ability to characterize the extent of depletion in specificities with differing prevalences. Overall, SIV-specific IgG was depleted to a similar extent in the antigen-specific as total plasma IgG (**Figure 5**): IdeS demonstrated a greater degree of IgG depletion as compared to Roz treatment, at approximately ten- and two-fold, respectively. While the degree of depletion varied among animals for each intervention, similar degrees of reduction were apparent across all antigens within a given animal, suggesting limited bias across specificities. Similarly, the kinetic profile of both depletion and recovery phases were consistent across antigens but varied between animals. SIV-specific IgG returned to similar levels post treatment as were observed at pre-treatment timepoints. In contrast, antigen-specific IgM and IgA were relatively unaffected by either treatment (**Supplemental Figure 4**).

**Figure 5.**
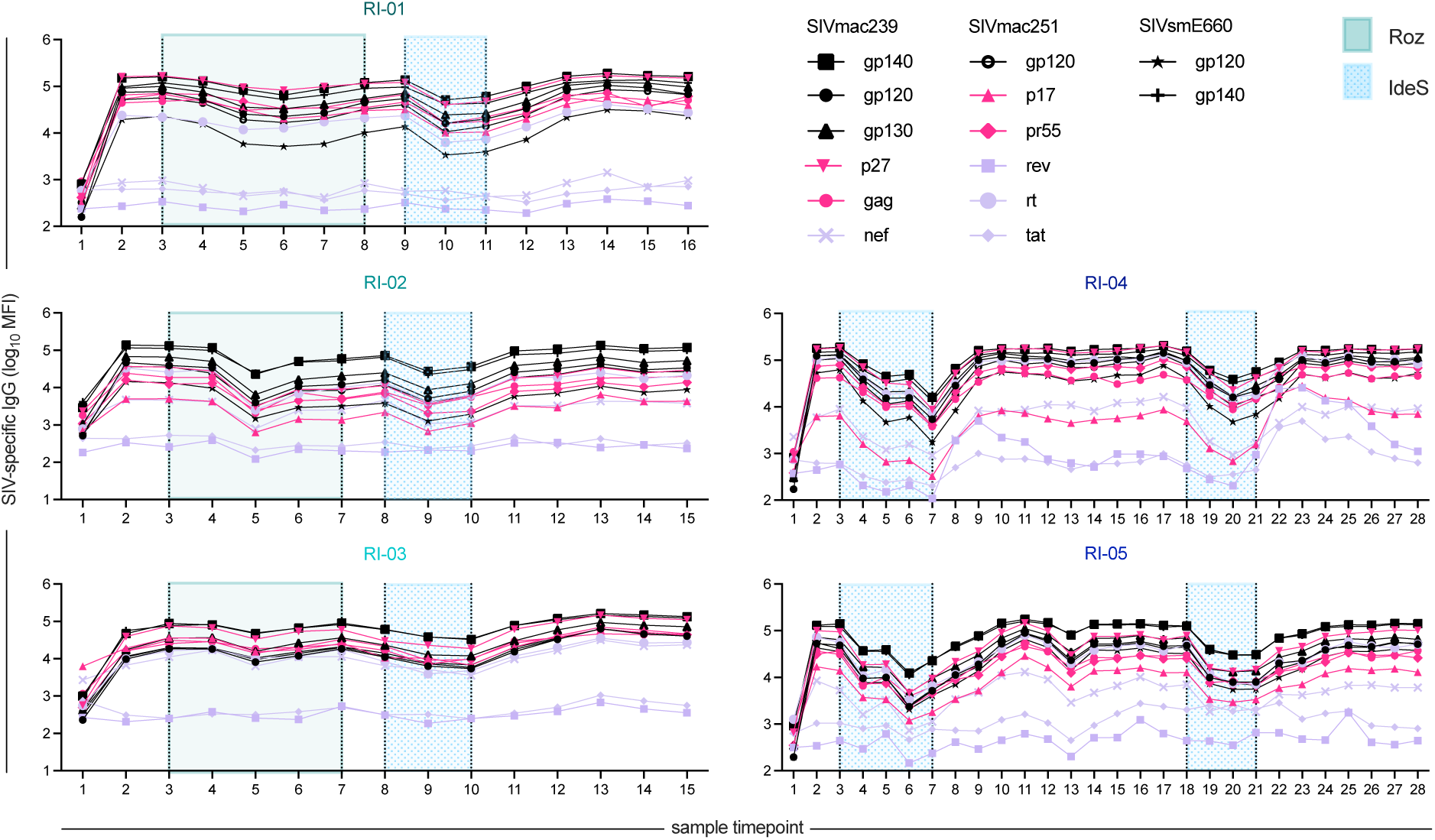
Levels of antigen-specific IgG over time in Roz- and IdeS-treated animals. Median fluorescent intensity (MFI) of serum IgG specific for a panel of SIV antigens over time in each of five treated animals. Animals were SIV-naïve at the first sample timepoint. Antigen specificity is indicated by shape and color with envelope, internal, and non-envelope structural proteins in black, purple, and pink, respectively. Sampling timepoints during Roz treatment are indicated in green, and IdeS treatment in blue.

### Alteration of humoral response post intervention

Within the SIV-specific IgG pool, levels of each IgG subclass were also assessed (**Figure 6**). In general, Roz treatment resulted in less effective depletion of each IgG subclass than did IdeS as detected by multiplex assay. Kinetics of depletion across animals and even between specificities within an animal varied in some cases. For example, capsid-specific IgG3 was not depleted by Roz treatment in animal RI-03, though it was in RI-02, and the rate of decay between p17- and p27-specific IgG3 in RI-01 were distinct. While only a subset of antigens induced an IgG3 response, varying from one to seven, typically including p27 (5/5 animals) and envelope (4/5 animals), all of these IgG3 specificities were thoroughly depleted by IdeS treatment. In contrast pigtail IgG2 was depleted efficiently by both Roz and IdeS, despite the incomplete cleavage of rhesus IgG2 in vitro.

**Figure 6.**
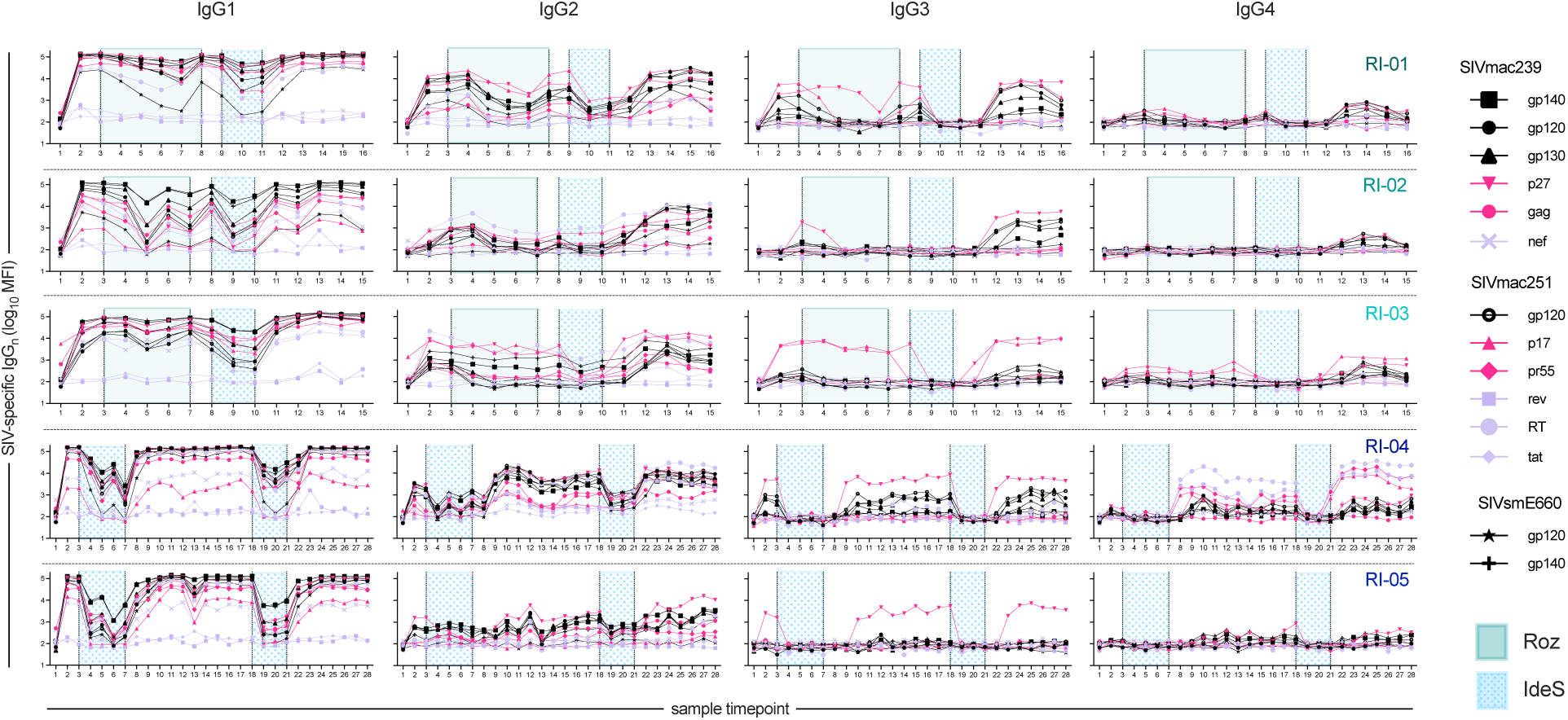
Levels of antigen-specific IgG subclasses over time in Roz- and IdeS-treated animals. Median fluorescent intensity (MFI) of SIV-specific IgG1, IgG2, IgG3, IgG4 (columns) detected in serum in each of five treated animals (rows) over time. Antigen specificity is indicated by shape and color with envelope, internal, and non-envelope structural proteins in black, purple, and pink, respectively. Sampling timepoints during Roz treatment are indicated in green, and IdeS treatment in blue.

Lastly, in contrast to IgG1, which exhibited similar levels and specificities before and after depletion, most animals showed some evidence of change in the level or presence of IgG2, IgG3, or IgG4 following depletion. Levels of IgG2, particularly to envelope, tended to increase after depletion. Increases in envelope-specific IgG3 were observed in animal RI-02. Most notable is animal RI-04, who exhibited almost a 100- fold increase in IgG4 responses to internal proteins rt and tat, as well as to structural proteins p17 and pr55.

### Functional consequences of IgG depletion

Given the incomplete cleavage of IgG by IdeS and the possibility that scIgG was detected by secondary reagents used to quantify isotypes and subclasses, we next wished to evaluate the impact of each IgG depletion strategy on the ability of antigen-specific antibodies to bind to FcyR. SIV-specific antibodies were detected with tetramerized rhesus Fcγ and FcαR (**Figure 7**). Here again, depending on the antigen-specificity, animal, and Fc receptor, different patterns of depletion emerged. In general, Roz had a more limited effect of detection of FcγR-binding antibodies, particularly in animals RI-01 and RI-03. For several animals, IdeS treatment resulted in ∼1000-fold signal reductions and the complete absence of detectable FcyR-binding SIV-specific antibodies. This effect was particularly pronounced for the low and moderate affinity FcyR (FcyRIIa, FcyRIIb, FcyRIIIa). In contrast, the high affinity FcyRI still showed signal above the pre-infection baseline sample for most animals, although high background observed for other antigens limited testing to envelope-specificities for this receptor. FcRn-binding antibodies showed an intermediate phenotype. FcαR-binding antibodies were more sporadically induced among animals and their levels varied within and outside of Roz and IdeS treatment periods.

**Figure 7.**
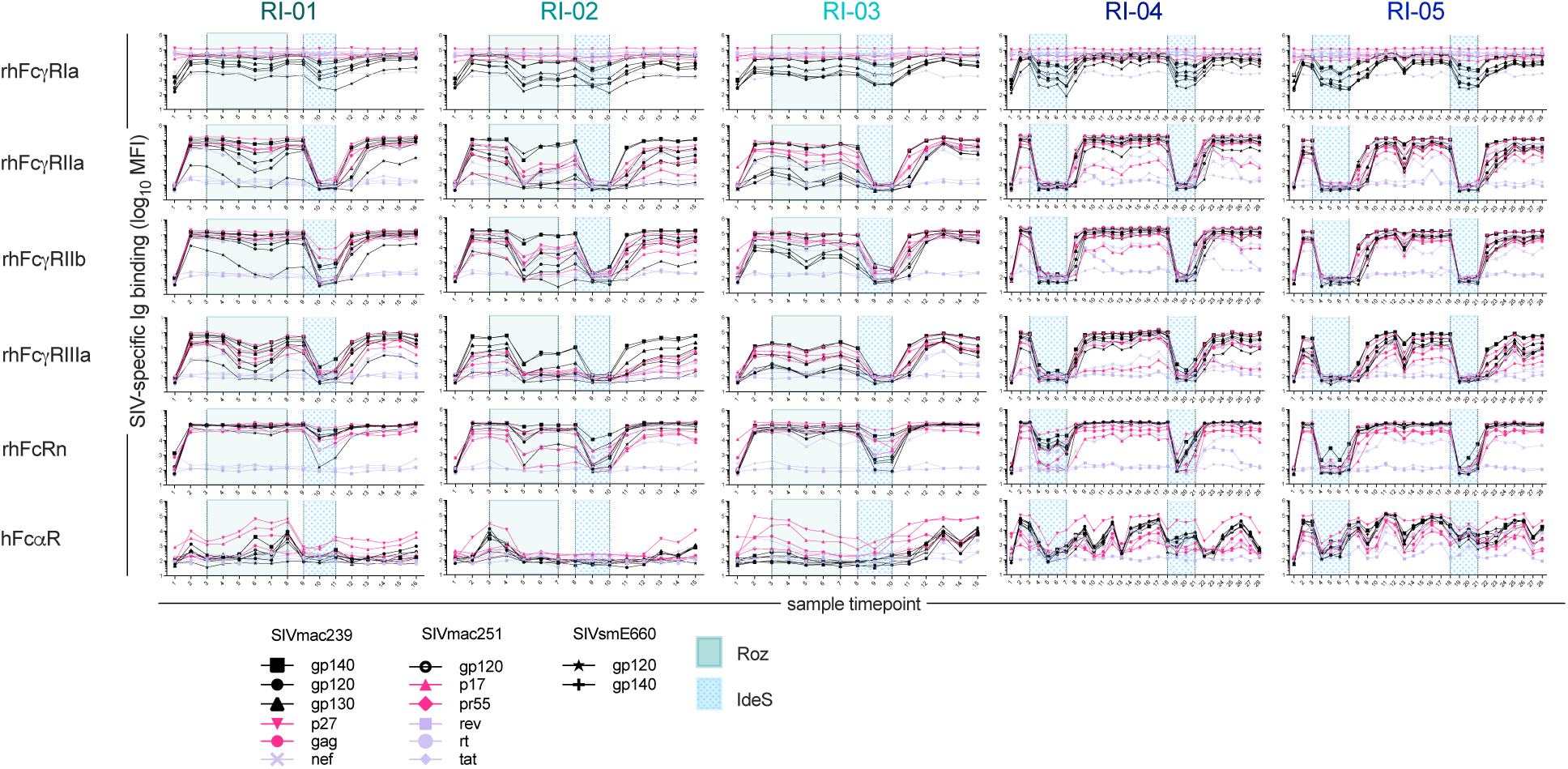
Levels of Fc receptor binding to antigen-specific antibodies over time in Roz-and IdeS-treated animals. Median fluorescent intensity (MFI) of rhesus FcR (rows) detected in serum in each of five treated animals (columns) over time. Antigen specificity is indicated by shape and color with envelope, internal, and non-envelope structural proteins in black, purple, and pink, respectively. Sampling timepoints during Roz treatment are indicated in green, and IdeS treatment in blue.

Lastly, the functional impact of Roz and IdeS treatment on the activity of SIV-specific antibodies was evaluated using a multiplexed assay of antibody-dependent complement cascade component C3b deposition (ADCD) (**Figure 8**). Roz treatment exhibited limited ability to reduce this activity, whereas in contrast, IdeS treatment reduced ADCD activity to baseline for multiple antigen-specificities in most of the animals.

**Figure 8.**
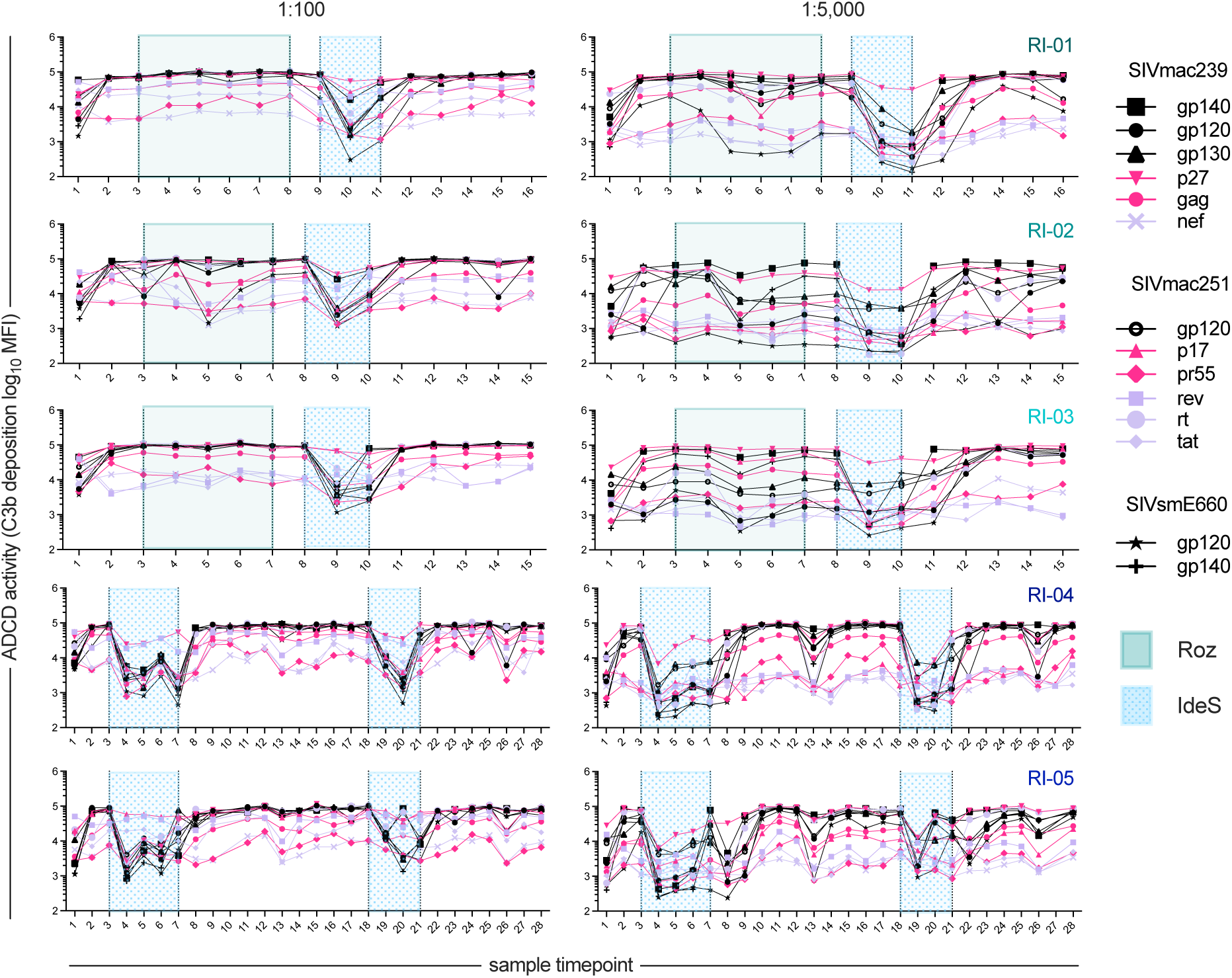
Antibody-dependent complement deposition (C3b) over time in Roz- and IdeS-treated animals. Median fluorescent intensity (MFI) of C3b deposition detected in serum for two serum dilutions (columns) in each of five treated animals (rows) over time. Antigen specificity is indicated by shape and color with envelope, internal, and non-envelope structural proteins in black, purple, and pink, respectively. Sampling timepoints during Roz treatment are indicated in green, and IdeS treatment in blue.

### IdeS is more efficient at depletion than Roz

Overall, among the five animals that received a total of seven IdeS treatment regimens, and the three animals that each received one Roz regimen, IdeS resulted in a greater degree of depletion (**Figure 9A**), which was achieved more rapidly but was not necessarily more durable. Extent of depletion of total IgG reflected comparable reductions in each of the antigen-specific subpopulations tested, though not necessarily in the detection of FcyR -binding antibodies or the ADCD function, which was at times undetectable despite the presence of residual antigen-specific IgG. Overall, using the regimens tested, IdeS treatment resulted in more effective IgG depletion in pigtail macaques.

**Figure 9.**
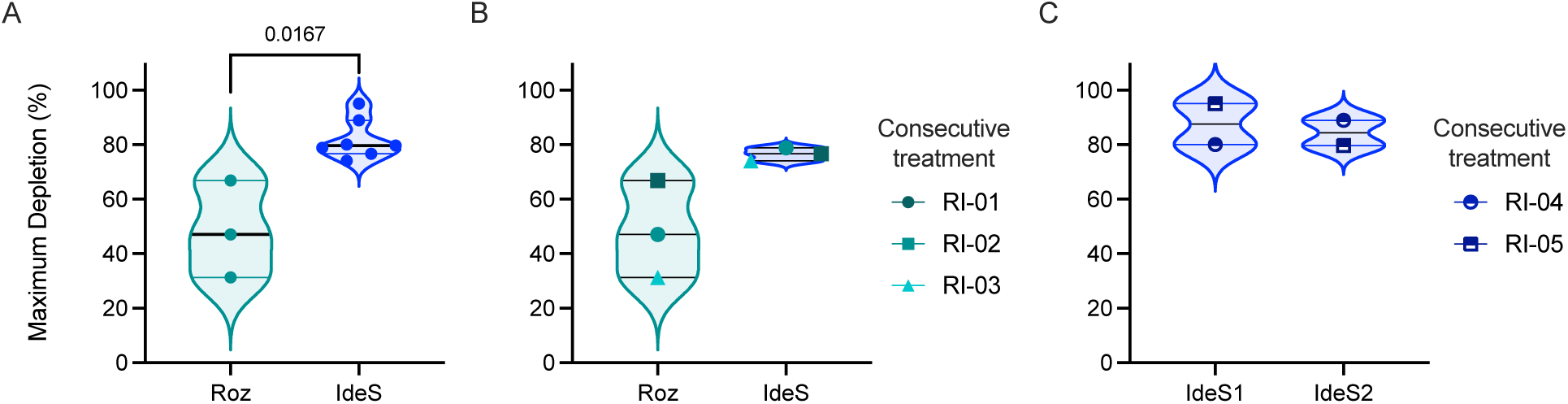
Comparison of maximal IgG depletion efficacy in Roz- and IdeS-treated animals. **A-C.** Maximum IgG depletion for each regimen of Roz and IdeS in all animals (**A**), in dually-treated animals (**B**), and in serially IdeS-teated animals (**C**). Statistical significance defined by Mann-Whitney U test.

Animals RI-01, RI-02 and RI-03, who underwent consecutive treatment with Roz then IdeS, showed different depletion efficiency. Roz depletion (Mean 48.4%, SD± 17.8%) was not as effective as the IdeS depletion (Mean 76.5%, SD± 2.5%) (**Figure 9B**). While some reduction in circulating IgG due to prior administration of Roz may contribute to the effect observed, each animal demonstrated a greater degree of maximum reduction of circulating IgG following IdeS treatment. Only animal RI-02 exhibited a similar degree of depletion for both interventions, suggesting the greater efficiency of IdeS in the context of matched animals. Lastly, animals RI-04 and RI-05 both underwent two rounds of IdeS with a depletion efficacy (Mean 87.6%, SD 10.6%, N=2) and (Mean 84.4%, SD 6.57%, N=2) (**Figure 9C**). Similar depletion efficiency and a lack of detectable anti-drug antibodies (**Supplemental Figure 5**) demonstrate the feasibility of repeated IdeS regimens.

## Discussion

The current study shows the high degree of conservation of the neonatal Fc receptor (FcRn) across humans, cynomolgus macaques, rhesus macaques, and pigtail macaques. Specifically, the FCGRT gene encoding the FcRn α-chain is found as a single copy in all these species, and no copy number variations have been documented^68^, underscoring the evolutionary stability of this immune regulatory mechanism. Importantly, while molecular analyses have identified minor sequence variants in the macaque FCGRT α-chain, such as S26N, G255S, A313S, and S355L, these sequence differences do not appear to result in meaningful changes to FcRn expression or function^69^. The protein structure of FcRn, including the approximately 365 amino acid α-chain, remains largely conserved between human and non-human primates, facilitating comparable mechanisms for recycling and transport of IgG^68^.

FcRn antagonists such as Efgartigimod^28^, Nipocalimab^29^, Batoclimab^23^, Orilanolimab^18,20^, and Rozanolixizumab^30–33^ are antibody-based therapies used clinically to treat various autoimmune conditions. In humans, IgG depletion efficiency by FcRn blockade differs amongst these treatments, though all typically report at least a 50% reduction [single dose Efgartigimod (50-85%)^28,35,70^, Nipocalimab (75-80%)^29,70,71^, Batoclimab (70-80%), Orilanolimab (50-70%), Rozanolixizumab (50-80%)^30,70^]. Duration of depletion varies as well, with Efgartigimod and Batoclimab reducing IgG levels for up to three months, followed by a two month period for Nipocalimab, and one month for Rozanolixizumab or Orilanolimab. Whereas Nipocalimab and Batoclimab have been associated with reductions in serum albumin, Efgartigimod^72,73^, Orilanolimab^73^, and Rozanolixizumab^73^ appear to spare albumin, a property with important implications for long-term tolerability and therapeutic index.

All of these FcRn-directed treatments effectively reduce serum IgG in cynomolgus macaques^35,61,74,75^, which are a frequently used model for immunoglobulin pharmacodynamics and pharmacokinetics. Beyond preclinical testing of these interventions for safety and efficacy, some have also been used to probe the role of antibodies in different disease states. For example, Roz has previously been tested with a single dose of 30 mg/kg or 60 mg/kg in rhesus macaques to measure the impact on donor-specific alloantibodies in a sensitized model and resulted in a 68% depletion of donor-specific IgG by four days post infusion^61^. Together, these findings underscore the relevance of FcRn antagonism as a therapeutic modality while highlighting pharmacodynamic heterogeneity among currently available drug candidates.

Here, in pigtail macaques chronically infected with SIV, we investigated two approaches to deplete IgG and determined their efficacy in reducing IgGs specific for several viral antigens. In rhesus macaques, 30 mg/kg of Roz given i.v. showed mean maximal IgG depletion at 68% (SD 10.6%) of baseline with recovery starting at seven days post treatment with full recovery by day 28^61^. This dose, followed by seven daily 5 mg/kg s.c. doses, increased depletion to 80%^66^. Our experiment in pigtail macaques, in which four weekly doses of Roz were administered i.v. at 30 mg/kg, showed a mean maximal depletion of IgG of 48.4% (SD 17.8%). In prior studies, anti-drug antibodies (ADA) developed in a majority of cynomolgus monkeys treated daily with Roz^31,33^. One limitation of our study is that we did not assess development of ADA to Roz. The observation that systemic IgG levels recovered to baseline by the fourth dose in one animal and almost to baseline by the third dose in another raises the possibility that ADA responses to Roz occurred.

In contrast to Roz, IdeS has been shown to rapidly cleave IgG, and its short half-life (4.9 ± 2.8 hours in humans at a dose of 0.24 mg/kg^47,76^) appears to be associated with lower risk of ADA in animal models^21^. In rhesus macaques, IdeS administered in a single i.v. dose of 2 mg/kg reduced circulating IgG by 70-80% with recovery to baseline by day 14^21^. Plasma IgG levels were reduced by ∼75-95% with a similar rate of recovery. As in a prior study^21^, ADA responses to IdeS were not observed.

While both IgG depletion methods were specific for IgG with no effect on IgM or IgA, Roz and IdeS are typically employed to investigate the effects of depleting antigen-specific fractions of the systemic antibody repertoire. The effect of these interventions on total serum Ig may or may not reflect an impact on antigen-specific Ig subsets. Notably, animals treated in our study were SIV-infected, affording the opportunity to monitor the depletion of antibodies specific to a panel of SIV proteins typically targeted by humoral responses during SIV infection. To this end, each treatment led to similar reductions of circulating total IgG binding across the 14 antigen specificities evaluated. Recovery rates of total IgG across antigen specificities were also similar, suggesting that both depletion and rebound phases were consistent across different specificities. Further, within antigen-specific fractions, levels of individual IgG subclasses could be tracked over time. Dominating the SIV-specific antibody fraction, IgG1 showed a similar degree of depletion as total antigen-specific IgG. However, the large dynamic range of detection of antigen-specific antibodies indicated that for some specificities, reductions by multiple orders of magnitude were observed. For less prevalent specificities, as well as for less common subclasses (e.g. IgG2, IgG3, IgG4), levels fell below the limit of detection in several animals. Similarly, the ability of antigen-specific antibodies to be bound by low affinity (IIa, IIb, IIIa) but not of the high affinity (I) FcyR was also often eliminated, consistent with substantial reduction in complement deposition activity observed in a functional assay. Collectively, this data suggests that the antigen-specific pools of total serum IgG are similarly depleted and that reductions in antibody function may match or exceed the extent of depletion observed among total serum IgG.

Though limited or no bias was observed in the efficiency of depletion of different specificities or IgG subclasses, several examples of differing kinetics or characteristics of rebounding antigen-specific antibodies were observed. For example, in some animals, antigen-specific IgG3 or IgG4 were not present before depletion but were observed post-treatment. While we cannot exclude that these differences are mediated by SIV infection rather than IgG depletion, changes in antibody profiles following therapy are of potential clinical relevance. Interestingly, little work has been reported to characterize the profile of antibody rebound post-treatment with these or other interventions clinically. While most studies look at the reduction of pathogenic antibodies and their recovery^28,61,77^ there are recent studies of Efgartigimod in myasthenia gravis demonstrating that, in addition to reducing serum IgG, treatment leads to the expansion of memory B cells and plasma cells, highlighting complex immunomodulatory effects beyond simple IgG removal^78^. In pemphigus, Efgartigimod-induced IgG depletion was also associated with the disappearance of desmoglein-specific B cells, indicating that anti-FcRn therapies may influence autoreactive B cell populations and potentially foster regulatory plasma cells with immunosuppressive properties^79^.

While this study suggests that both Roz and IdeS can be useful tools in pigtail macaque models of disease, alternative methods for depletion of pathogenic antibodies as well as means to interfere with their activities also exist. These include high dose intravenous immunoglobulin (IVIg), leading to saturation of the FcRn receptor and lower levels of pathogenic IgG, or therapeutic plasma exchange, involving the physical removal of circulating plasma and replacement with donor plasma or albumin, thereby directly eliminating pathogenic antibodies and immune complexes from the bloodstream ^36,37^. Additionally, blockade of FcyR (FcyRIIa^80^ and FcyRIIIa^81^) is being investigated as a complementary approach for reducing pathogenic activity without directly influencing the levels of pathogenic antibodies. This menu of strategies, which can also be employed in combination, presents opportunities to balance maintenance of the activities of beneficial IgG while reducing activity of pathogenic species.

There are several limitations to this study. Effects of IdeS and Roz on the B cell compartment, albumin, or antibody in tissues were not evaluated. In addition, our study groups included small numbers of animals: three were treated with Roz and then received IdeS, while two underwent two rounds of IdeS. Levels of antigen-specific antibody binding to a diversity of Fc receptors was tested, but only one antibody effector function assay was evaluated *in vitro*. Lastly, all animals were infected with SIV, which provided the opportunity to examine antigen-specific antibody depletion and rebound, and one antibody mediated cellular effector function (ADCD), but may have influenced the degree and kinetics of these processes in association with changes in viral latency and tissue or systemic viremia.

In summary, our study establishes pigtail macaques as a model for anti-FcRn and IdeS efficacy in IgG depletion and provides additional data on rebound of IgG subclasses and antigen-specific IgGs. Notably, IgG subclass levels did not always return to pre-depletion baselines; newly detected or altered levels of some IgG subclasses appeared after depletion. Loss of IgG led to a measurable reduction in FcyR binding and complement c3b deposition confirming altered IgG mediated functions. This finding suggests that analyzing IgG subclasses after intervention in clinical studies could be valuable, especially for autoimmune conditions influenced by specific pathogenic subclasses and their distributions.

## Data Availability Statement

Tabulated raw data from this study is available from the corresponding author by reasonable request.

## Acknowledgments

The Anti-FcRn [RozR1LALA] antibody (NHPRR CAT-00095, RRID:AB_2888630) and IdeS (NHPRR, CAT-00341) used in this study were provided by the Nonhuman Primate Reagent Resource (NHPRR, RRID:SCR_012986), funded by NIAID U24 AI126683.

The following reagents were provided by the NIH Nonhuman Primate Reagent Resource (NIAID U24 AI126683): Anti-rhesus IgG1 [ena]-biotin antibody, Anti-rhesus IgG2 [dio]-biotin antibody, Anti-rhesus IgG3 [tria]-biotin antibody, and the Anti-FcRn [RozR1LALA] antibody (Cat# PR-0001, RRID: AB_2888630) provided by the Nonhuman Primate Reagent Resource (NHPRR, RRID: SCR_012986). ChimeraX was developed by the Resource for Biocomputing, Visualization, and Informatics at the University of California, San Francisco, with support from National Institutes of Health R01-GM129325 and the Office of Cyber Infrastructure and Computational Biology, National Institute of Allergy and Infectious Diseases.

Further acknowledgements for other reagents are provided in Supplemental Table 3.

