## Supplemental Materials for "Depletion and recovery of IgG following treatment with Rozanoliximab and Imlifidase in pigtail macaques"

**Supplementary Materials**

| **Supplementary Figures** | |
|---|---|
| Supplementary Figure 1 | Serum protein levels over time in treated animals |
| Supplementary Figure 2 | Total IgA, IgM, IgG measurements in additional animals that underwent Roz treatment. |
| Supplementary Figure 3 | Total IgA, IgM, IgG measurements in additional animals that underwent IdeS treatment. |
| Supplementary Figure 4 | Levels of antigen-specific IgA and IgM over time in Roz- and IdeS-treated animals. |
| Supplementary Figure 5 | Levels of IdeS anti-drug IgG responses |

| **Supplementary Tables** | |
|---|---|
| Supplementary Table 1 | IgG alignment accession numbers |
| Supplementary Table 2 | Total immunoglobulin assay |
| Supplementary Table 3 | Fc array assay |

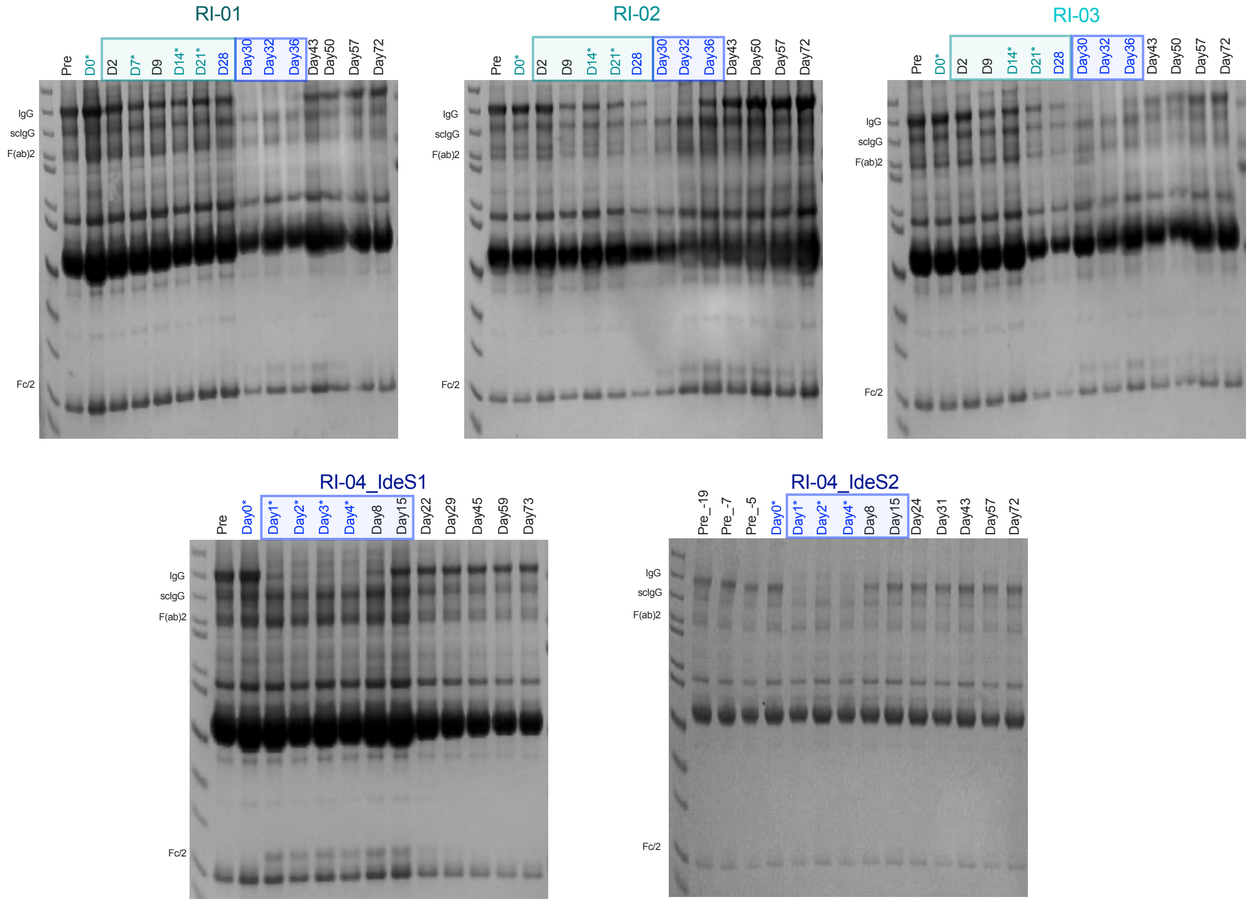


**Supplemental Figure 1: Serum protein levels over time in treated animals**. SDS-PAGE of total serum protein in each animal over time during Roz and/or IdeS treatment periods. Days of Roz or IdeS treatment are indicated with an asterisk and those anticipated to show IgG depletion in the teal and blue boxes, respectively.

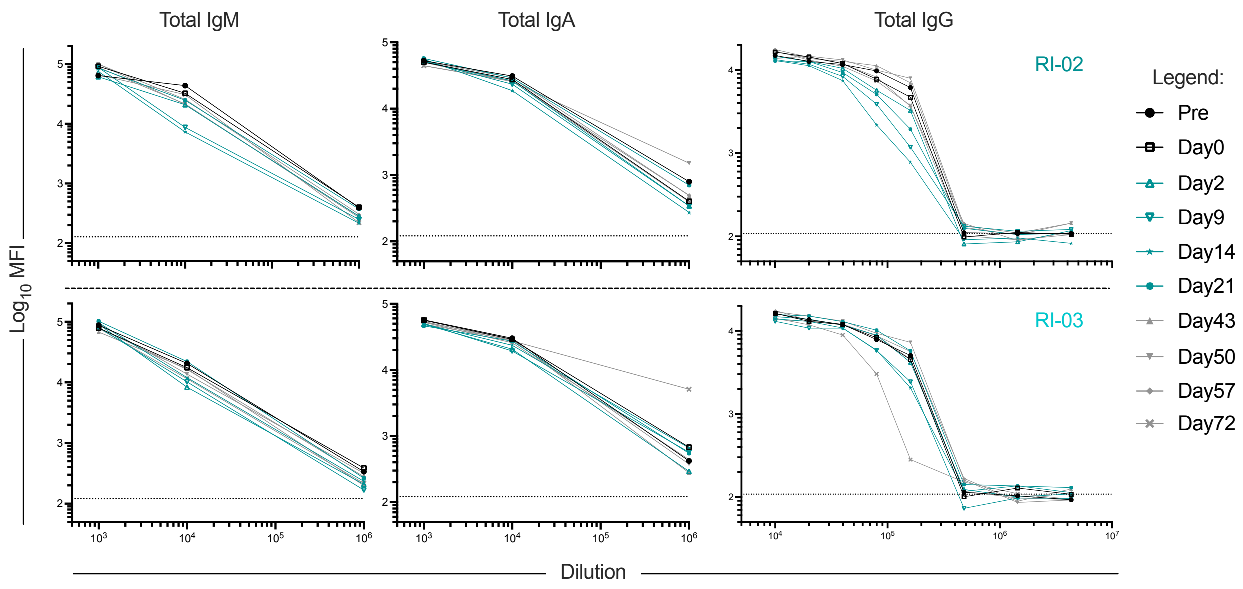


**Supplemental Figure 2: Total IgA, IgM, IgG measurements in additional animals that underwent Roz treatment**. Median fluorescent intensity (MFI) of total IgM (left), IgA (center), and IgG (right) measured in diluted serum over time in animals RI-02 and RI-03 (rows). Color and symbol indicate sample timepoint. Dotted line indicates lower limit of detection.

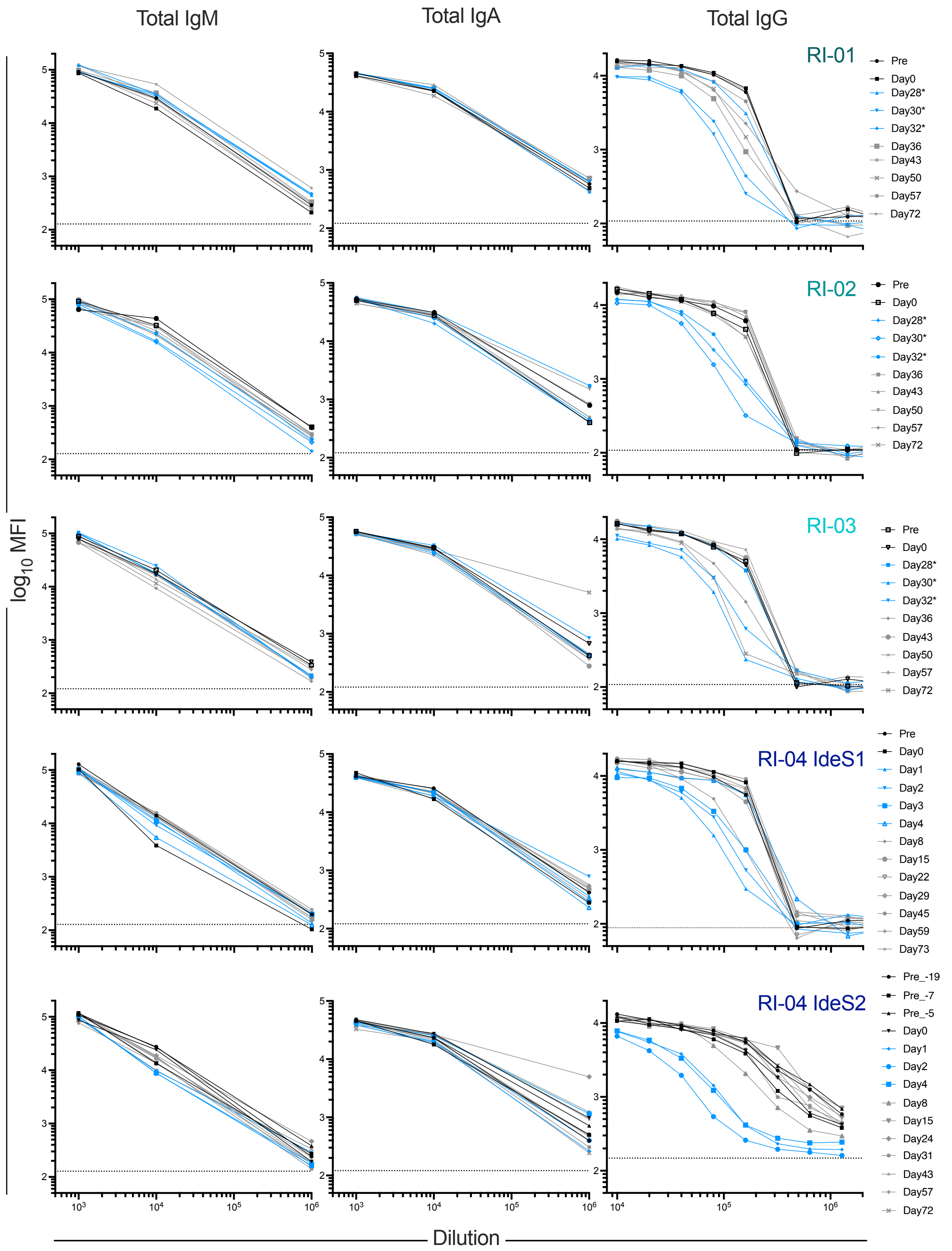


**Supplemental Figure 3: Total IgA, IgM, IgG measurements of the additional animals that underwent IdeS treatment.** Median fluorescent intensity (MFI) of total IgM (left), IgA (center), and IgG (right) measured in diluted serum over time in treated animals (rows). Color and symbol indicate sample timepoint. Dotted line indicates lower limit of detection. Color of the line denotes the timepoint with various dilution on X axis.

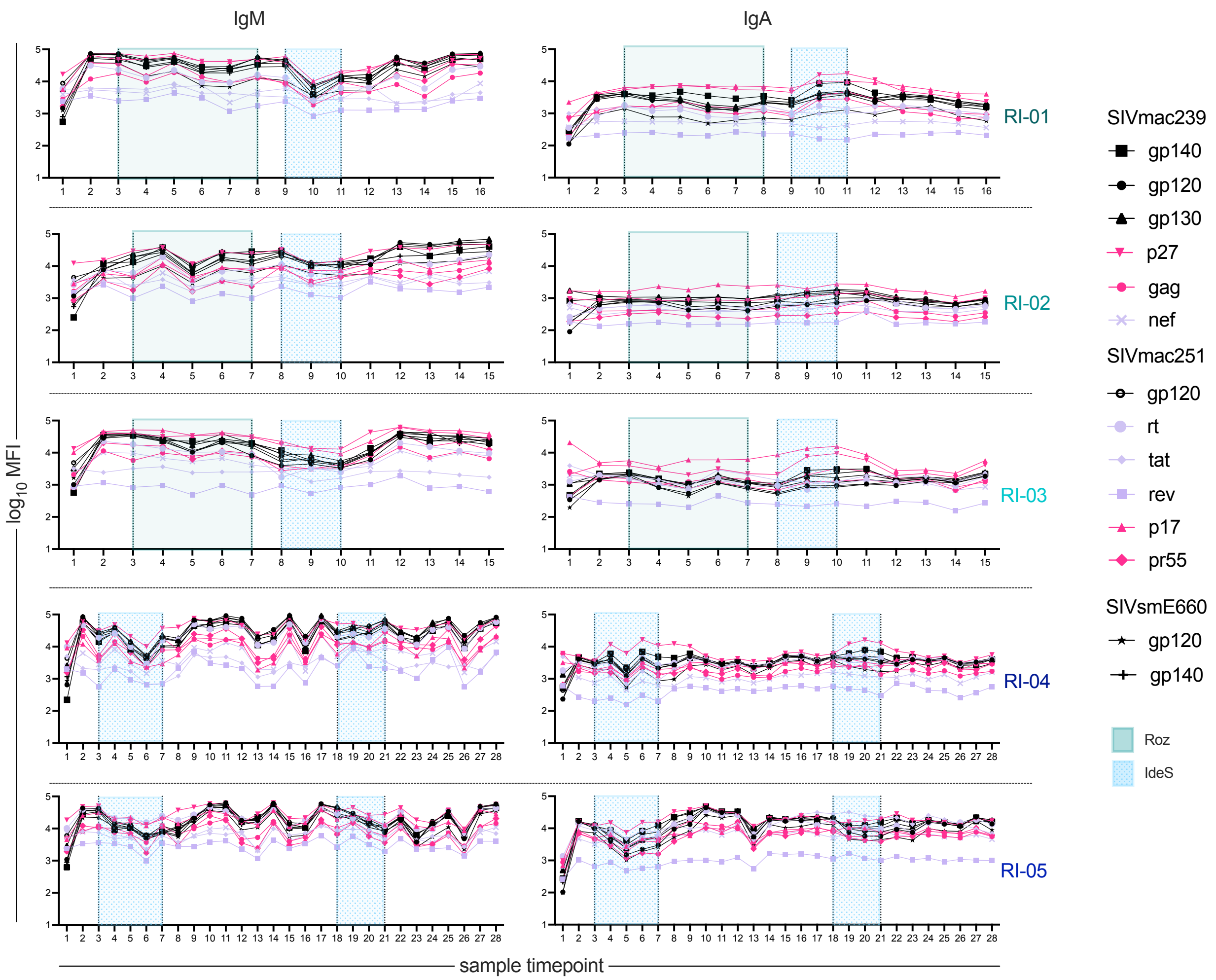


**Supplemental Figure 4: Levels of antigen-specific IgA and IgM over time in Roz and IdeS treated animals**. Median fluorescent intensity (MFI) of SIV-specific IgA (left) and IgM (right) (columns) detected in serum in each of five treated animals (rows) over time. Antigen specificity is indicated by shape and color with envelope, internal, and non-envelope structural proteins in black, purple, and pink, respectively. Sampling timepoints during Roz treatment are indicated in green, and IdeS treatment in blue.

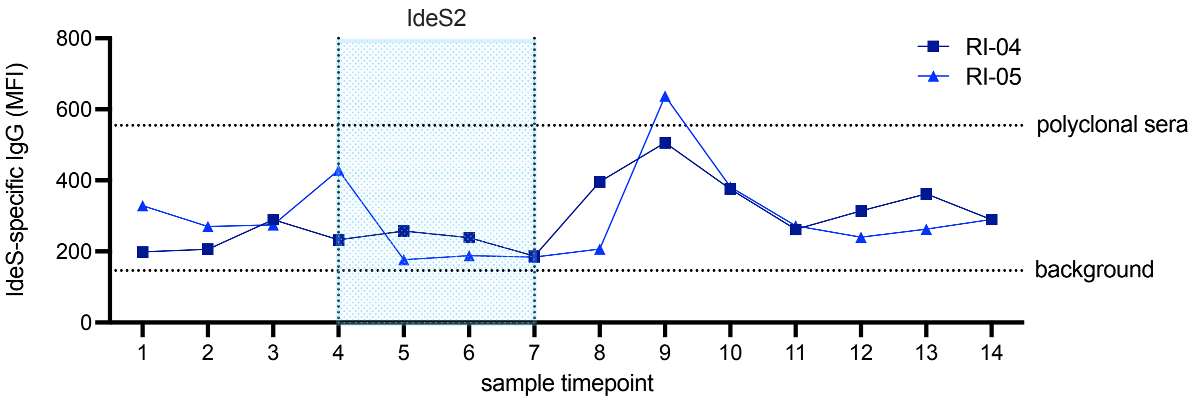


**Supplemental Figure 5: Measurement of anti-IdeS IgG in doubly treated animals RI-04 and RI-05.** Median fluorescent intensity (MFI) of IdeS-specific IgG before, during and after the second IdeS treatment in animals LJ02 and LJ04. Bottom dotted line indicates average of technical blanks. Top dotted line indicates average of two wells of a Rhesus polyclonal pool.

| Alignment accession numbers | | |
|---|---|---|
| **Species** | **Antibody** | **NCBI Numbers** |
| Human | IgG1 | J00228 |
| | IgG2 | J00230 |
| | IgG3 | D78345 |
| | IgG4 | K01316 |
| Rhesus | IgG1 | AF045537 |
| | IgG2 | AF045539 |
| | IgG3 | AF045538 |
| | IgG4 | AY29250 |
| Cynomolgus | IgG1 | JQ868728 |
| | IgG2 | JQ868729 |
| | IgG3 | JQ868730 |
| | IgG4 | JQ868731 |
| Pigtail | IgG1 | JQ868732 |
| | IgG2 | JQ868733 |
| | IgG3 | JQ868734 |
| | IgG4 | JQ868735 |

**Supplemental Table 1: Accession numbers used in the IgG alignment.**

| Total immunoglobulin assays | | |
|---|---|---|
| Fc Detection | Source | Serum dilutions tested |
| a-IgG | Southern Biotech 4700-09 | 1:10,000- 1:4,320,000 |
| a-IgM | Southern Biotech 9020-09 | 1:1000 - 1:1,000,000 |
| a-IgA | Southern Biotech 2050-09 | 1:1000 - 1:1,000,000 |

**Supplemental Table 2: Antibodies and dilutions used in the total immunoglobulin assays.**

| Detection Reagents | | | |
|---|---|---|---|
| Fc Detection | Source | Serum dilution tested | |
| a-IgG | Southern Biotech 4700-09 | 1:20,000 | |
| a-IgG1 | NHPRR Cat#PR-7116 | 1:10,000 | |
| a-IgG2 | NHPRR Cat#PR-0003 | 1:500 | |
| a-IgG3 | NHPRR Cat#PR-0006 | 1:100 | |
| a-IgG4 | NHPRR Cat#PR-7186 | 1:100 | |
| a-IgA | Southern Biotech 2050-09 | 1:150 | |
| a-IgM | Biotechne NBP1-73556 | 1:500 | |
| rhFcR1a | Chan et. al (PMID: 27559046) | 1:20,000 | |
| rhFcR2a | Chan et. al (PMID: 27559046) | 1:20,000 | |
| rhFcR2b | Chan et. al (PMID: 27559046) | 1:20,000 | |
| rhFcR3a | Chan et. al (PMID: 27559046) | 1:20,000 | |
| rhFcRN | Chan et. al (PMID: 27559046) | 1:500 | |
| huFcaR | Duke Protein Production Facility | 1:500 | |
| Antigens | | | |
| Antigen | Company | Cat # | Acknowledgement |
| gp120 SIVmac239 | Immune Technology | IT-001-022p | |
| gp120SIVmac251 | Immune Technology | IT-001-156p | |
| gp120 SIVsmE660 | Immune Technology | IT-001-142p | |
| gag SIV-1/mac239 | Immune Technology | IT-001-024EP | |
| Nef SIV-1/ mac239 | Immune Technology | IT-001-026Ep | |
| gp140 SIVmac239 | Immune Technology | IT-001-140p | |
| p27 SIVmac239 | Immune Technology | IT-001-018EP | |
| gp140 SIVsmE660 | Immune Technology | IT-001-141p | |
| SIVmac239 gp130 | BEI | ARP-12797 | Obtained through the NIH HIV Reagent Program, Division of AIDS, NIAID, NIH: Simian Immunodeficiency Virus (SIV) SIVmac239 gp130-His Recombinant Protein, ARP-12797, contributed by Dr. Klaus Uberla. |
| SIVmac251 pr55 gag | BEI | ARP-13384 | Obtained through the NIH HIV Reagent Program, Division of AIDS, NIAID, NIH: Simian Immunodeficiency Virus SIVmac251 pr55 Gag Recombinant Protein, ARP-13384, contributed by NIAID, DAIDS. |
| SIV p17 (SIVmac251) | NIBSC | 660 | Obtained through the Programme EVA Centre for AIDS Reagents, NIBSC, UK: SIV p17 (SIVmac251), NIBSC 660, contributed by Dr. R. Randall, University of St Andrews. |
| SIVmac251 clone J5 RT | NIBSC | 686 | Obtained through the Programme EVA Centre for AIDS Reagents, NIBSC, UK: SIVmac251 clone J5 RT protein, NIBSC 686, contributed by FIT Biotech Oyj Plc, Tampere, Finland. |
| SIVmac251 clone J5 rev | NIBSC | 684 | Obtained through the Programme EVA Centre for AIDS Reagents, NIBSC, UK: SIVmac251 clone J5 rev protein, NIBSC 684, contributed by FIT Biotech Oyj Plc, Tampere, Finland. |
| SIVmac251 clone J5 tat | NIBSC | 685 | Obtained through the Programme EVA Centre for AIDS Reagents, NIBSC, UK: SIVmac251 clone J5 tat protein, NIBSC 685, contributed by FIT Biotech Oyj Plc, Tampere, Finland |

**Supplemental Table 3: Materials used in SIV-specific immunoglobulin assays.**
